# Alleviation of arsenic toxicity in rice by employing silver nanoadsorbent: Insights into growth reconditioning with antioxidant defence

**DOI:** 10.64898/2025.12.07.692792

**Authors:** Triparna Mukherjee, Saptarshi Konar, Tapan Kumar Das

## Abstract

The present work imparts an efficient, simple and cost-effective nano-based technology for the removal of arsenic (III) from aqueous environment that was subsequently investigated on plant system for its evaluation. The study was designed to investigate the arsenic induced toxicity in rice cultivar MTU 7029 and its amelioration by monolayer protected silver nanoadsorbent under laboratory conditions. An efficient amelioration of arsenic induced toxicity was successfully achieved by the supplementation of nano-pretreated water (∼97% arsenic free) in hydroponic culture medium of rice seedlings (MTU 7029). As (III) exposure in the growth medium declined the seedlings growth, physiological conditions as well as it disrupted antioxidant defence system by overproducing reactive oxygen species (ROS) and superoxide anions. However, the application of the nanoadsorbent exhibited the alleviation of As (III) toxicity through the improvement of seedlings growth, restoration of normal root morphology and anatomy as indicated by scanning electron microscopy (SEM) and histology; decreased level of root oxidizability, electrolyte leakage, hydrogen peroxide (H_2_O_2_), malondialdehyde (MDA) and proline content. Furthermore, the supplementation of nano-pretreated water in hydroponic culture medium of the rice seedlings showed improved level of antioxidant enzymes such as, catalase (CAT), ascorbate peroxidase (APX), superoxide dismutase (SOD), glutathione reductase (GR) and reduced level of ROS generation and superoxide anions formation as revealed by CM-H2DCFDA and DHE staining of the roots. Hence, the present findings conveying a successful amelioration of arsenic induced toxicity in rice plant could be a useful technique for the alternative use of irrigation water in the arsenic contaminated land.

**Highlights:**

- Arsenic (As III) exposure led to reduced growth and oxidative damage in rice plant.
- Silver nanoadsorbent efficiently mitigated As (III) toxicity.
- Nanoadsorbent improved plant growth by enhancing the antioxidant defence system.
- The technique served a sustainable solution for rice cultivation in As contaminated land.

## 1. Introduction

Plants are highly flexible with their optimal atmospheres and tolerant to the various environmental stresses through different stress combating strategies for attaining better growth and their development (Majumder et al., 2019). Among the various stress components, heavy metals are one of the prime factors that affect severely the crop productivity globally. These metals that cause toxicity to plants are biologically non-essential in nature (Ghori et al., 2019). Arsenic is one of such non-essential metals categorized by the International Agency of Research on Cancer (IARC) as a class I toxicant and food chain pollutant that shows acute and also chronic toxicity according to the kind of its exposure (Zhao et al., 2010). Arsenic enters into the food chain primarily through the contaminated ground water or directly by the consumption of phyto-based food materials, like seeds, fruits and vegetables grown on arsenic affected soils (Finnegan and Chen, 2012) Arsenic accumulation in soil from groundwater is of great concern in agricultural field due to its adversative effects on growth of economically important crops (Shaibur 2021). Unfortunately, this polluted groundwater has often been utilized for irrigation purpose in various countries, that has eventually amplified the extent of As in the soil. The raised level of arsenic reduces the plant nutrient absorption efficiency, and hinders the plant growth and productivity (Shaibur et al., 2012). Among different cereal crops, Rice (*Oryza sativa* L.) is the second most important staple crop supplying over half of the world’s population (Hassan et al., 2022; Shaheen et al., 2022). Arsenic stress has a massive ill effect on rice productivity (Majumder et al. 2020; Upadhyay et al., 2020). The phytotoxic effects of arsenic on rice mainly depend on its oxidation state. Among the two different oxidation states, As may exist in the environment as both inorganic (arsenite and arsenate) and organic (monomethyl arsenic acid-MMA and dimethyl arsenic acid-DMA) forms (Chen et al. 2017). However, both the toxicity and availability of arsenite (AsIII) in rice plant are higher than arsenate (Kalita et al., 2018). The presence of As (III) can lead to interaction with hydrophilic groups of plant enzymes and proteins, resulting in loss of their functions and potential cell damage (Garg and Singla, 2011). While plants are exposed to As stress, they undergo competition to maintain a balance between toxicity and tolerance, that can eventually lead to the plant death (Finnegan and Chen, 2012). The phytotoxicity of As caused oxidative stress in rice plants through enhancing the reactive oxygen species (ROS) generation, negatively effecting defense system of plants, and impairing cell membrane damage caused by lipid peroxidation (Kushwaha et al., 2019; Nahar et al., 2022). Apart from these, plants also modulate a series of enzymatic and non-enzymatic antioxidant defense mechanisms to combat such As induced oxidative stress. Plants respond to the oxidative stress by modulating different antioxidant enzymes, like superoxide dismutase (SOD), catalase (CAT), ascorbate peroxidase (APX), Glutathione reductase (GR) etc. Despite the enzymatic protection system, plants also retain a broad range of various molecules that are able to scavenge free radical, such as ascorbate, glutathione, flavonoids, carotenoid, anthocyanins and vitamins etc. (Hossain et al., 2006). These molecules perform the role of singlet and triplet oxygen quenchers as well as the decomposer of peroxide (Hossain et al., 2012).

Several studies have been carried out on oxidative stress and defense mechanism in plants under As stress (Shri et al., 2009: Li et al., 2019; Altowayti et al., 2022). Various agricultural techniques have been employed to minimize the accumulation and toxicity of As in rice plants, such as irrigation practices, supplementation of minerals as fertilizer, nanotechnological applications along with biotechnological approaches (Jalil et al., 2023). Very recently, the implementation of nanotechnology in field crops such as rice has confronted advancements in agricultural production worldwide (Jiang et al., 2021; Farooq et al., 2022; Khan et al., 2025). Nanotechnology deals with the formation of nanoparticles (NPs) with typical dimensions ranging from 1 to 100 nm, having a unique physiochemical and structural characteristic (Khan et al., 2019). Among different metal-based NPs, silver (Ag) nanoparticles have now gained attention in agricultural practices. In this context, the present study was designed by considering a new approach towards the protection against As induced toxicity in rice plant using an ecofriendly nontoxic nano adsorbent. In the present study, Arsenic contaminated water was pretreated in presence of the synthesized silver nanoparticle by the process of adsorption and finally at the end, the nano adsorbents were separated from water by ultracentrifugation, for the safe use of nano-adsorbents. In this study, following the pretreatment procedure, it has been targeted to assess the reduced arsenic toxicity in the rice plant. Hence, the present study was aimed to investigate the arsenic induced toxicity and its amelioration by nano-absorbents in rice plant by studying the morphological and anatomical changes, the degree of oxidative damages and other different biochemical attributes.

## 2. Materials and methods

### 2.1. Chemicals

All chemicals and solvents used in the present study were of superior ultrapure grade. The Hoagland nutrient medium, glutaraldehyde, isoamyl alcohol, sulphosalicylic acid of HiMedia, India; mercuric chloride, hydrogen peroxide, nitric acid, hydrochloric acid, sulfuric acid, phosphoric acid, glacial acetic acid, sodium hydroxide, sodium borohydride, absolute alcohol, formaldehyde and ninhdrin of Merck, India; sodium arsenite, 2’,7’-dichlorofluorescin diacetate, dihydroethidium, thiobarbituric acid, bovine serum albumin, proline standard of Sigma-Aldrich, USA; potassium iodide, trichloroacetic acid, di sodium EDTA, dithiotreitol, polyvinyl pyrrolidone, phenylmethylsulfonyl fluoride, triton X-100, L-methionine, riboflavin, nitro blue tetrazolium, N,N,N,N-tetramethyl ethylenediamine, oxidized glutathione, nicotinamide adenine dinucleotide phosphate reduced tetrasodium salt, ascorbic acid, Acrylamide, Bis-acrylamide, potassium ferricyanide, ferric chloride, 3-[4,5-dimethylthiazol-2-yl]-2,5-diphenyltetrazolium bromide and 2,6-Dichlorophenolindophenol (DCPIP) of Sisco Research Laboratory (SRL), India, were used throughout the experiments.

### 2.2. Source of Plant material, growth conditions and treatment

Viable rice seeds of selected cultivar MTU 7029 were procured from Rice Research Station, Chinsurah, West Bengal. Seeds were first surface sterilized by using 0.1% (w/v) mercuric chloride (HgCl_2_) for 5min followed by repeated washing with sterile deionized water. Sterile seeds were then germinated on moistened filter papers in sterile petri-dishes for 48hrs under dark condition, afterward shifted under white fluorescent light (52µEm^-2^s^-1^ PAR) at 25°C for three more days for proper germination and growth. Subsequently, depending on the emergence of the radical, uniformly germinated seeds were chosen and transferred to grow hydroponically in autoclavable plastic jars containing modified 1/4^th^ strength of Hoagland nutrient medium (Hoagland and Arnon, 1950) and the medium supplemented with 10, 25, 50 and 100 µM sodium arsenite (NaAsO_2_), as well as with its respective treatment dose and nanoparticle control at pH 6.2 for next 10days under regulated environment in growth chamber with 16hr photoperiod at 25°C with 60-70% relative humidity (RH). The concentrations of arsenic selected in our study were based on our initial experiments where LD_50_As was recorded as 200 µM (Data not shown). During the entire experiment the nutrient medium was replaced with fresh one after every two days.

For the purpose of this study monolayer protected silver nanoparticles, Ag@MSA in a background of aqueous system get involved in the interaction with arsenic As (III) through batch mode procedure. The nanoadsorbent used in the present study was previously synthesized following the methods of our earlier report by Mukherjee et al. (2019). This synthesized silver nanoparticle (Ag@MSA) in addition with its interaction study with As (III) was performed as a continuation of our previous work as well. During the interaction study a 1:5 ratio of Ag@MSA /As (III) concentration was maintained and the process was continued till the equilibrium reached. Here a constant stock concentration of arsenic As (III) (from where the toxic dose of arsenic was added to the medium for arsenite mediated toxicity study) was used for the interaction with a constant dose of Ag@MSA that was also previously standardized in our previous study by Mukherjee et al. (2019). A stock concentration of 500 μM (37.45 mg/L As concentration) of As (III) solution was used for this interaction study. After the completion of the interaction the adsorbent was separated from the solution by ultra-centrifugation at 60,000 rpm for 1:30 hours at 4°C. This supernatant water was taken for the entire experiment for the uptake by rice plant (MTU 7029) as a beneficial alternative of arsenic contaminated toxic water.

### 2.3. Experimental setup

Experiment was conducted on 15 days old seedlings of rice cultivar MTU 7029, grown hydroponically on 1/4^th^ strength of Hoagland nutrient medium were served as control (CON), while sodium arsenite solution and nano absorbents based pretreated water supplemented hydroponic Hoagland medium were assigned as As and TR respectively. A positive control (AgNP) was also taken for each experimental setup. Hence, total ten sets of treatment combination (CON, As10, As25, As50, As100, / TR10, TR25, TR 50, TR100 and AgNP) were used for each experiment in triplicate, arranging the total setup in fully randomized manner.

### 2.4. Harvest procedure

15days old seedlings were harvested followed by proper washing with milli-Q water and exploited for the study of several parameters. Roots and leaves were separated and weight individually for further experimentations. After harvesting, experiments were divided into three sets on the basis of which the samples were preserved. First set for arsenic content analysis where fresh sample was used just after harvesting. In second set also the freshly collected samples were used for the measurement of root & shoot length, detection of ROS accumulation and cell death, histochemical as well as histological and SEM analyses. The third set samples were frozen in liquid nitrogen, immediately after harvesting and stored at -80°C for biochemical analyses.

### 2.5. Arsenic content analysis

Plant parts both root and leaf tissues were selected for arsenic content analysis. Tissues were properly washed with deionized water and dried in an oven at 45-50°C till the samples become dried. Then the samples were weighted and crushed to get a powder form. For determining the initial arsenic content in the seeds, the seed samples were also considered in the present study. In case of seeds the samples were directly weighted and crushed, without any drying procedure. Thereafter, for the acid digestion 2mL of concentrated HNO_3_ and 1mL of H_2_O_2_ were added to the 100mg of samples (dried form) and kept for overnight. Following the digestion, the samples were filtered through a Milli-pore membrane (0.45 mm; CASL 45 2.5CMD, membrane: acetyl cellulose) and shifted quantitatively to a volumetric flask and made up with Milli-Q water. The samples were retained in a plastic container for assessment. The standard reference material of arsenic (E-Merck, Germany) was applied for the calibration as well as quality assurance purpose for every analytical batch the standard references as well as blank solution were also processed under the similar conditions. The arsenic content was quantified by using Hydride Generation-Atomic Absorption Spectrometry [Varian Model AA 140 (USA)] method (HG-AAS). A solution of 1.25% sodium borohydride in 0.5% sodium hydroxide and 5 M HCl were utilized for hydride generation purpose. Rice flour 1568a (National Bureau of Standards, Gaithersburg, MD, USA) was used as a Standard Reference Material (SRM) for the present study. Detection limit of arsenic was 0.003 mg/L.

### 2.6. Morphologial and anatomical studies

Root and shoot length of control (CON), positive control (AgNP), arsenic affected (As) and treated (TR) were measured in centimeter (cm) followed by anatomical studies, immediately after harvesting on 15^th^ day. For study of anatomical alteration, the roots were fixed in FAA (95% ethanol: glacial acetic acid: formaldehyde: water, 10:1:2:7), dehydrated through a graded ethanol series (20, 30, 50, 70, and 90%) sequentially for 20 min at each step with three changes and finally in 100% ethanol for 30 min. Then the samples were embedded in paraffin wax (Merck, Germany) the thin sections (10μm) were cut using an electronic rotary microtome (MICROMHM 340E, Thermo Fisher Scientific, Germany) followed by staining with Hematoxylin and Eosin stain. The section strips were further washed with xylol, mounted with DPX (BDH) and examined under a compound microscope (Leica, DME, Germany).

### 2.7. Scanning electron microscopy: sample preparation and imaging

Root samples (root tips and rest of the parts of CON, As, TR and AgNP) for scanning electron microscopy were fixed in 2.5% glutaraldehyde in 0.1 M phosphate buffer (pH 7.0) for 4 h at 4^°^C and dehydrated through a graded ethanol series (30%, 50%, 70%, 90% and 100%) sequentially for 20 min each with two changes. Then the root samples were passed through a sequential series of absolute alcohol and isoamyl alcohol (3:1, 2:2, 1:3) for 30 min in each. Finally, tissues were kept in isoamyl alcohol for 30 min. The tissue was then dried in a critical point dryer (HCP-2, Hitachi, Japan) and coated with gold sputter prior to the examination under a scanning electron microscope (ZEISS EVO LS 10, Germany).

### 2.8. ROS imaging with CM-H2DCFDA and DHE staining

In case of ROS imaging, root tips of CON, As, TR and AgNP were stained with 12.5 μmol mL^-1^ of CM-H2DCFDA (chloromethyl derivative of 2’,7’-dichlorofluorescin diacetate) for 10 min and then washed with Phosphate buffer saline. The fluorescence images of the stained roots were observed using a fluorescence microscope (Zeiss, AxioCam MRm, Scope.A1, excitation 400–490 nm, emission 520 nm). Likely, ROS formation was examined following the procedure of Yamamoto et al. (2002) using a superoxide anion (O^2-^) specific indicator, dihydroethidium (DHE). Following the excision of root tips, staining was done with 10 µM DHE in 100 mM CaCl_2_ (pH 4.7) for about 30 min. Prior to the observation of stained roots under fluorescence microscope (Zeiss, AxioCam MRm, Scope.A1, excitation 488 nm, emission ≥ 505 nm), the roots were soaked in 100 mM CaCl_2_ for 10 min for the removal of residual dye.

### 2.9. Electrolyte Leakage (EL)

Electrolyte leakage (EL) was estimated by measuring the ions leaching from the tissue into deionized water (Dionisio-Sese and Tobita, 1998). Fresh samples of both root and leaves (100mg) of CON, As, TR and AgNP were cut into small pieces and incubated in deionized water at 25°C for duration of 2 hr in test tubes. Initial conductivity (E1) of the bathing solution was recorded with an electrical conductivity meter. The tubes containing both root and leaf samples were then boiled for 30 minutes to release all electrolytes and kept at 25°C to cool before recording the final conductivity (E2). The EL was calculated as a percentage data using the formula, EL (%) = (E1 / E2) × 100

### 2.10. Root oxidizabilty (RO)

Root oxidizability was determined to measure the roots’ oxidizing ability in terms of red triphenyl formazan formed by TTC reduction assay (Batish et al., 2007). 100mg of root tissues of CON, As, TR and AgNP were mixed with 5 mL of TTC (0.4% w/v) and 5 mL of 50mM phosphate buffer (pH 7.0). The mixture was then incubated for 3hr at 40° C. After addition of 2mL of 2N H_2_SO_4_, the roots were homogenized in 10mL of ethyl acetate to extract formazan. The absorbance of the extracted material was measured at 485 nm and RO was expressed as A_485_ g^-1^h^-1^.

### 2.11. Hydrogen peroxide and superoxide anion estimation

Hydrogen peroxide (H_2_O_2_) content was analyzed as per the method of Velikova et al. (2000). 100 mg of fresh root samples of CON, As, TR and AgNP were homogenized in ice bath with 0.1% (w/v) trichloroacetic acid (TCA) and centrifuged at 12, 000 g for 15 min. Then 0.5 mL of the supernatant was added into the 1 mL of 1 M potassium iodide (KI) and 0.5 mL of 50 mM potassium phosphate buffer (pH 7.0). The absorbance of the mixture was recorded at 390 nm. H_2_O_2_ content was determined using the extinction coefficient 0.28 µM^-1^ cm^-1^ and expressed as µM g^-1^ FW.

Superoxide anion activity i.e the rate of O^2-^ production was measured by observing the oxidation of epinephrine to adrenochrome according to the method of Misra and Fridovich (1971). The root samples (∼50mg) of CON, As, TR and AgNP were cut into small segments and placed in 2mL of reaction mixture having 100 µM di sodium EDTA, 20 µM NADH and 20 mM sodium phosphate buffer (pH 7.8). The reaction was initiated with the addition of 100 µl of 25.2 mM epinephrine (freshly prepared in 0.1N HCl). The samples were then shaken at 150 rpm for 5 min at 28°C on a rotary shaker. The tissues were then removed and the change in absorbance was measured at 480nm over a period of 5 min. The amount of adrenochrome produced was estimated using the extinction coefficient of 4.0 × 10^3^ µM^-1^ and expressed as nM g^-1^ FW.

### 2.12. Measurement of Proline and malondialdehyde (MDA) content

Proline quantification was done spectrophotometrically using the method of Bates et al. (1973). Frozen leaf tissues (100 mg) of CON, As, TR and AgNP was homogenized with 5 mL of 3 % sulphosalicylic acid and centrifuged at 12 000 g for 10 min. In test tubes 1mL of supernatants were mixed with 1mL of glacial acetic acid and 1 mL acid-ninhdrin [acid-ninhdrin was prepared by heating 1.25 g ninhydrin in 30 mL of glacial acetic acid and 20 mL 6 M phosphoric acid, with constant stirring, until dissolved. By cooling this reagent, it can be stored at 4°C for 24 hours] and boiled for 1 hour at 100°C. The reaction was then terminated in an ice bath. Reaction mixture was extracted with 2 mL toluene, stirred vigorously and kept at room temperature for 30 min till the separation of the two phases. The upper phase (1 mL), choromophore comprising toluene portion was warmed to normal room temperature and then the optical density was measured at 520 nm by using toluene as blank. Proline content was estimated using a standard curve of Proline standard and expressed as µM g^-1^ FW.

Malondialdehyde (MDA) is known as the ultimate decomposition outcome of lipid peroxidation and has been treated as an indication for lipid peroxidation status. Concentration of MDA, a major TBARS (Thiobarbituric acid reactive substances), was measured following the procedure of Hodges et al. (1999). Frozen leaf tissues (100 mg) of CON, As, TR and AgNP was grounded in 80% cold ethanol and centrifuged at 10,000 rpm for 5 min to pellet the debris. 1mL of supernatant in separate test tubes was added with 1 mL of 20% (w/v) trichloroacetic acid (TCA) containing 0.5% (w/v) thiobarbituric acid (TBA) or with 1 mL of 20% (w/v) TCA. Both the combinations were allowed to react in a system of water bath at 95° C for 1hr. Thereafter, samples were cooled in ice bath and centrifuged at 8000g for a duration of 10 min. Absorbance of the supernatant was recorded at 532 nm compared to a blank and there was subtraction from the value found at 600 nm for the correction of unspecific turbidity. The MDA content was expressed in terms of nmol gm^-1^ F.W. by using an extinction coefficient value of 155 mM ^-1^ cm^-1^.

### 2.13. Assay of antioxidant enzymes

Frozen root and leaf tissues (50 mg) of CON, As, TR and AgNP were homogenized in 50 mM potassium phosphate buffer (pH 7.8) comprising with 1 mM dithiotreitol (DTT), 1% (w/v) polyvinyl pyrrolidone (PVP), 1mM EDTA, 1 mM PMSF using ice-cold mortar and pestle kept at 4°C. Thereafter the centrifugation of the homogenate was done at 15,000g at 4°C for 30 min and the clear supernatant was utilized for enzymatic assays, such as catalase (CAT), Ascorbate peroxidase (APX), Superoxide dismutase (SOD) and Glutathione reductase (GR). Estimation of soluble protein was also carried out according to Bradford (1976) using bovine serum albumin (BSA) as the standard. All spectrophotometric studies were performed using an UV/visible Spectrophotometer (Genesis 10S UV-VIS, Thermo Scientific).

The assay for catalase activity was followed by Aebi (1984). For the analysis of catalase (EC 1.11.1.6) activity, suitable aliquot of enzyme was added into 50 mM potassium phosphate buffer (pH 7) and 20 mM H_2_O_2_ with a final volume of 1mL. The disappearance of H_2_O_2_ was observed by evaluating the decrease of absorbance at 240 nm for 3min at 30sec interval. With the extinction co-efficient value (∈=0.0436 mM^-1^ cm^-1^), the catalase activity was presented as mM H_2_O_2_ min^-1^ mg^-1^ protein. For peroxidase analysis, the activity of ascorbate peroxidase (APX, EC.1.11.1.7) was quantified through the oxidation of ascorbate to dehydroascorbate, as explained by Nakano and Asada (1981). In this assay, the frequency of hydrogen peroxide dependent oxidation of ascorbate was measured in a reaction combination containing 50 mM potassium phosphate buffer (pH 7.0), 0.5 mM ascorbate, 0.1 mM EDTA, 0.1 mM H_2_O_2_ and 0.1 mL of enzyme extract. Following the oxidation of ascorbate, there was a decrease of absorbance at 290 nm (∈= 2.8 mM^-1^ cm^-1^) for 3min at 30sec interval. The activity of APX was presented as μmol of ascorbate oxidized min^-1^ mg^-1^ protein.

Superoxide dismutase (SOD) (EC 1.15.1.1) activity was measured by nitro blue tetrazolium (NBT) photochemical analysis procedure (Beyer and Fridovich,1987). In this process, 1mL final volume of reaction mixture containing 50 mM potassium phosphate buffer (pH 7.8), 57 µM of NBT, 9.9 mM L-methionine and 0.025% (v/v) triton X-100 was added in 10 mL glass vial followed by the addition of 0.02 mL of sample (enzyme extract). The reaction was initiated with the addition of 0.01 mL of riboflavin solution (4.4 mg/100 mL) as well as subsequent placement of the vials in a black paper lined box with 20-W (Philips 20 W) fluorescent lamps for a period of 7 min. In case of positive control, sample volume was replaced by buffer. After completion of the illumination, absorbance of the reaction mixture solution was observed at 560 nm. A negative control (non-irradiated complete reaction mixture) was served as a blank. SOD activity was presented as U mg^-1^ protein. One unit of SOD was the amount of protein needed to inhibit 50% primary reduction of NBT under light.

The activity of glutathione reductase (GR) (EC 1.6.4.2) was measured through observation of the glutathione dependent oxidation of NADPH (Carlberg and Mannervik, 1985). 1mL final volume of reaction mixture containing 0.2 M of potassium phosphate buffer (pH 7), 2 mM NADPH, 2 mM EDTA and 20 mM oxidized glutathione (GSSG) was started to react followed by the addition of 0.1 mL of enzyme extract. The decline in absorbance (GSSG) value at 340 nm was observed for about 3min at 30sec interval. The activity of glutathione reductase was analyzed by using the extinction coefficient value for NADPH (6.2 mM^-1^cm^-1^) and presented as nmol NADPH became oxidized min^-1^ mg^-1^ protein.

### 2.14. Statistical analysis

Statistical analysis was performed using the data obtained independently from three repeated experiments. The whole experiment was carried out in a randomized block design. Results were analyzed as mean values ± standard errors and significant difference at (*p* ≤ *0.05*) among mean values were assessed by analysis of variance (ANOVA) and Duncan’s multiple range test (DMRT) using the MSTAT-C software program (ver.1.41, Michigan State University).

## 3. Results and discussion

### 3.1. Differential accumulation of As content in leaf and root

From the initial studies it was observed that approximately 97% of arsenic As (III) was removed due to the interaction of surface carboxylate group of monolayer protected silver nanoparticles (Ag@MSA) with As (III) within a time of 60 minutes at 30°C and neutral pH. During the study on rice seedlings (MTU 7029) arsenic uptake increased in both the leaves and roots with increasing As (III) concentrations in the hydroponic growth medium. Primary arsenic content in seed was recorded as 0.736889 μg As g-1 from AAS data. Notably, after exposure the arsenic accumulation was much higher in roots compared to leaves (Table 1). Rice seedlings treated with exogenously applied As (III) accumulated maximum As content at As100 µM in both leaves and roots; however, the seedlings supplemented with nano-pretreated (TR) water in hydroponic culture medium exhibited reduced accumulation of As (III) as compared the respective As concentrations tested alone (Table 1). The positive control AgNP resembling with CON seedlings in As accumulation directed a preliminary confirmation of As non-toxicity to the rice seedlings. The present result accounting to significantly higher accumulation of As in roots was also in accordance with the previous findings by many authors in different plant systems (Rahman et al., 2015; Koley et al., 2023). Roots are the primary site of interaction with different toxic elements for its’ higher accumulating nature as suggested by many researchers in their earlier reports (Su et al., 2010).

**Table 1.**
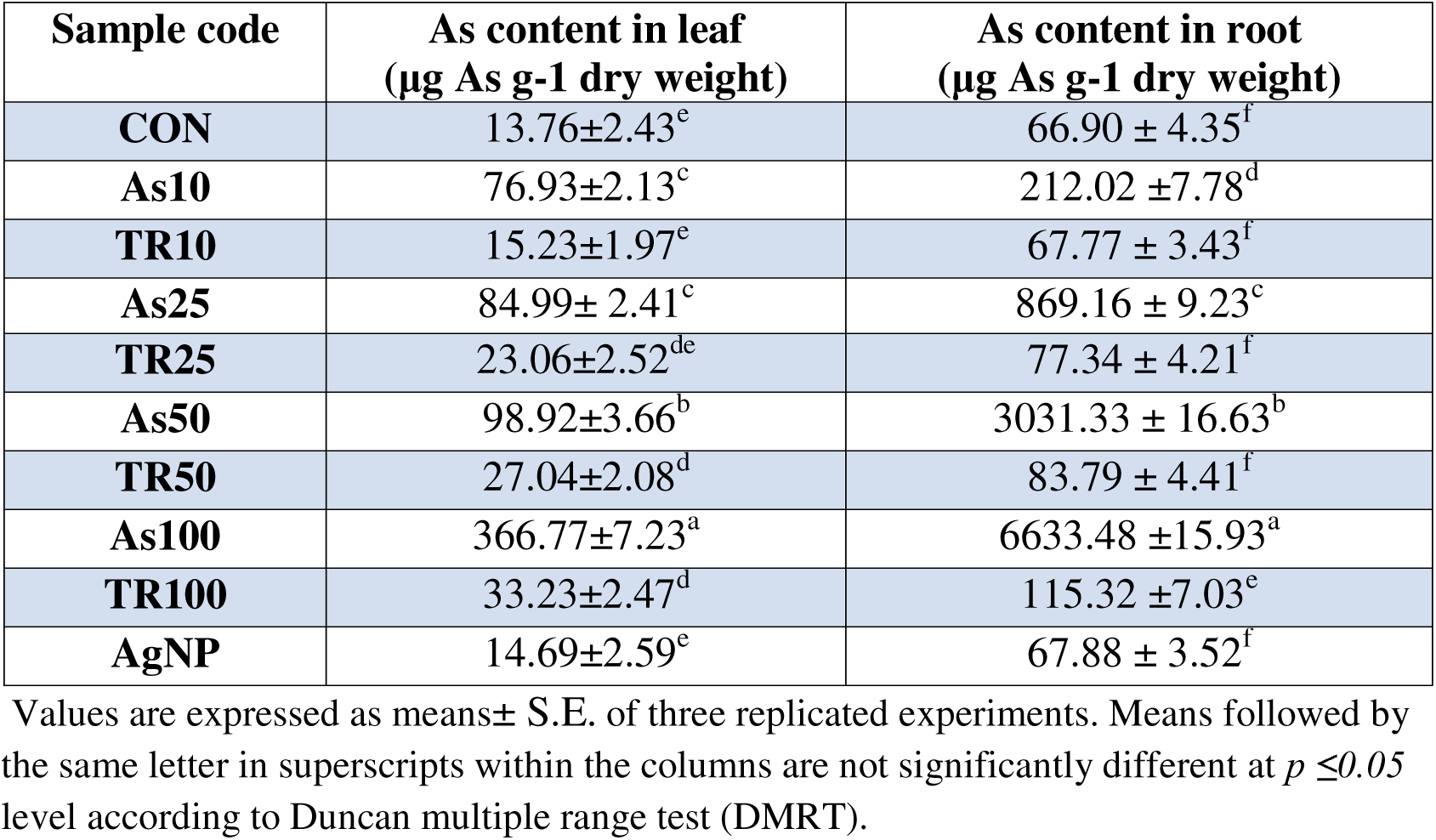
As content of 15days old seedlings of rice cultivar MTU 7029, grown hydroponically on 1/4^th^ strength of Hoagland nutrient medium.

### 3.2. Effect of As affected and nano-pretreated (TR) water supplementation on seedling growth and root anatomy

Rice seedlings (MTU 7029) growth were severely affected upon the exposure of different concentrations of As (III) used in the present study (Fig.1). This resulted in a significant reduction on root and shoot lengths that was recorded after harvesting on 15th day (Fig. 2). The macroscopic morphological alterations included not only the inhibition of root and shoot lengths, but also decreasing numbers of lateral roots and the main root apex looked necrotic and slimy with the increasing As concentration. The root length was decreased significantly in 50 and 100 µM of As doses which were more than 60% of reduction, whereas the reduction in shoot length was comparatively lower accounting to nearly 28% reduction, on an average. This inhibitory effect of As on root and shoot growth is evident with the earlier reports in many plant species (Armendariz et al., 2016; Singh et al., 2018). Such inhibitory effect was more pronounced in roots compared to shoots, as also accordance with previous studies on rice plant (Nath et al., 2014). To combat this As induced toxicity in seedlings, various strategies have been taken by several workers (Rahman et al., 2015; Kandhol et al., 2022). However, nanoparticle mediated alleviation of such arsenic induced toxicity has been merely reported in plant species (koley et al., 2023; Jalil at al., 2023). In the present work, seedlings grown on nano-pretreated (TR) water supplemented hydroponic medium showed the significant ameliorative effect with the increase of root, shoot length and formation of root hairs as similar to the control. Positive control AgNP exhibited nomal root and shoot growth that indicated no toxicity of AgNP to the rice seedlings.

**Fig. 1.**
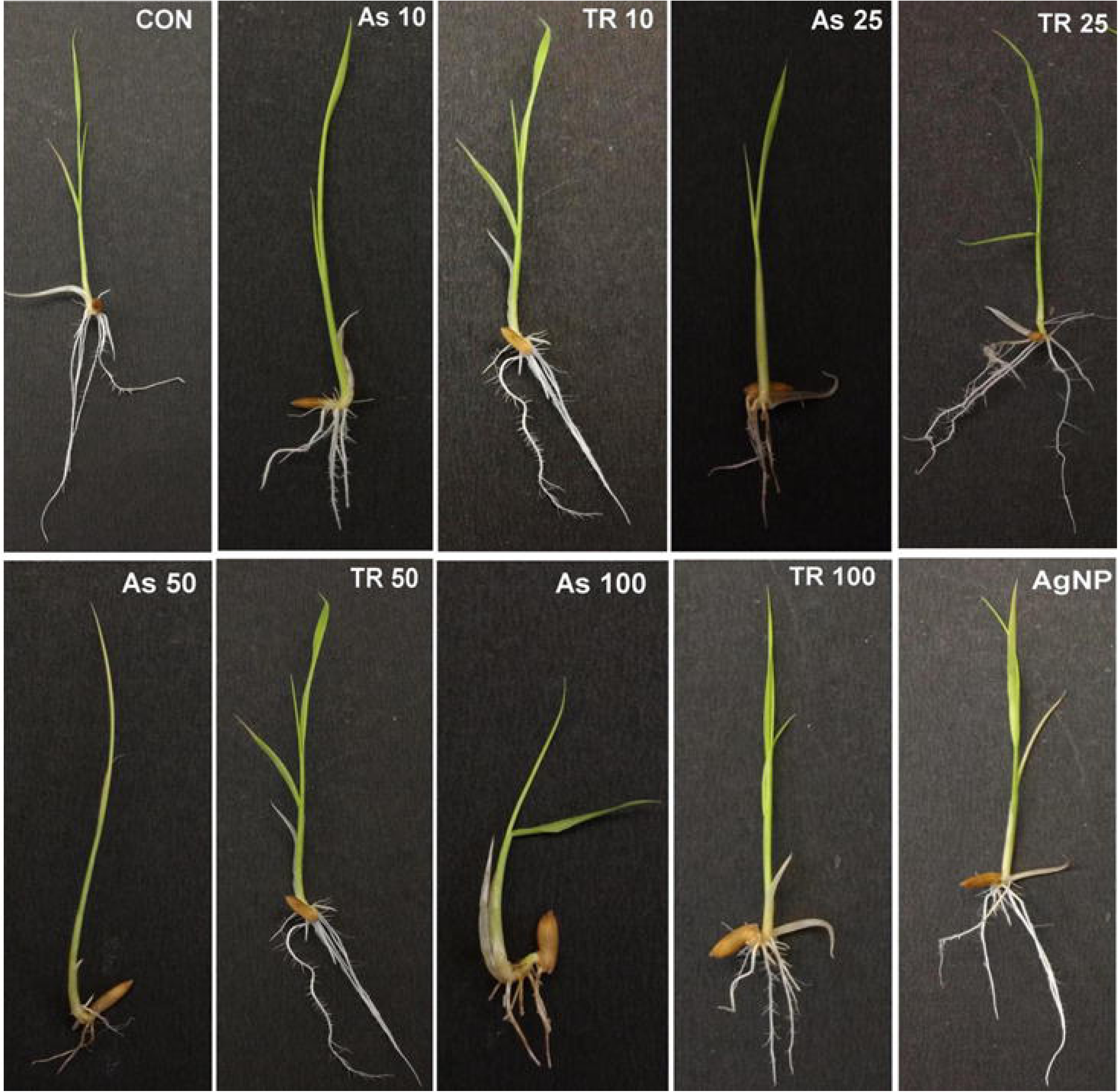
Morphological macroscopic photographs of 15days old rice seedlings grown on hydroponic nutrient medium amended with different concentrations of As(III),TR and AgNP.

**Fig. 2.**
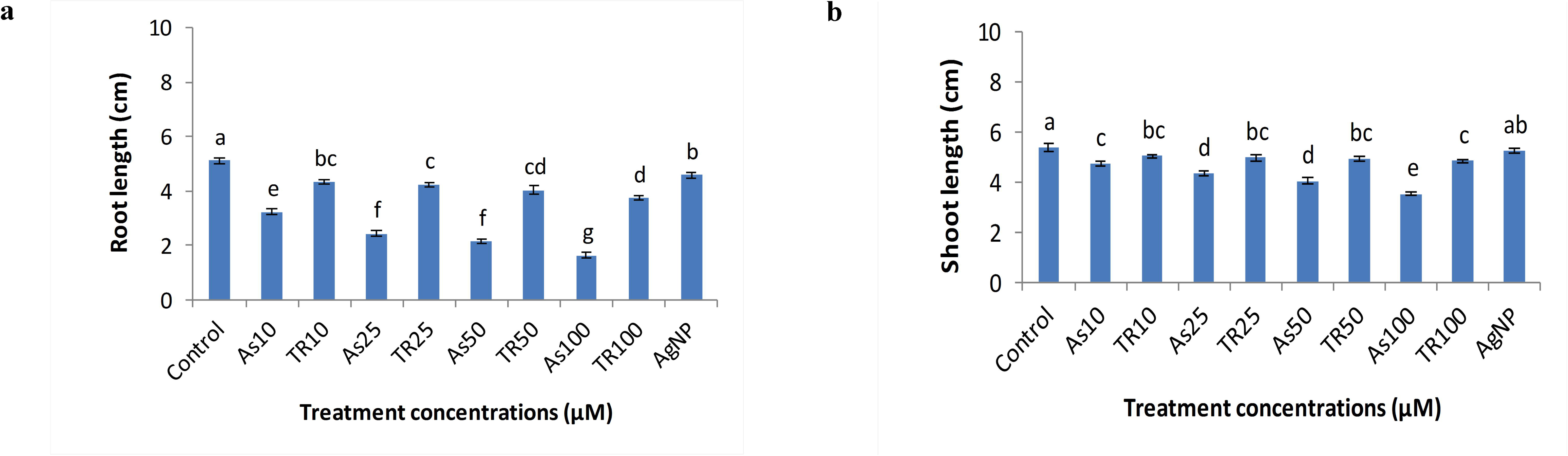
Response of rice seedlings to different concentrations of As (III), TR and AgNP in terms of root length (a) and shoot length(b). Data are expressed as means± S.E. of three replicated experiments. Vertical bars indicate standard errors. Mean values in columns followed by the different letters are significantly different at p ≤0.05 level according to Duncan multiple range test (DMRT).

Arsenic exposure also affected the root anatomy which was evident from the earlier studies on different plants upon uptake of various heavy metals by roots (Armendariz et al., 2016). In the present work, the epidermal cells were intact and the cortex region was normal in shape with uniseriate pericycle made up of thin-walled parenchyma cells in both the CON and AgNP, but in the As exposed root epidermal layer and the arenchymatous cortex region were distorted. Whereas in all respective treated (TR) concentrations, the damaging effect due to As exposure was almost nullified and somehow looked similar to control (Fig. 3).

**Fig. 3.**
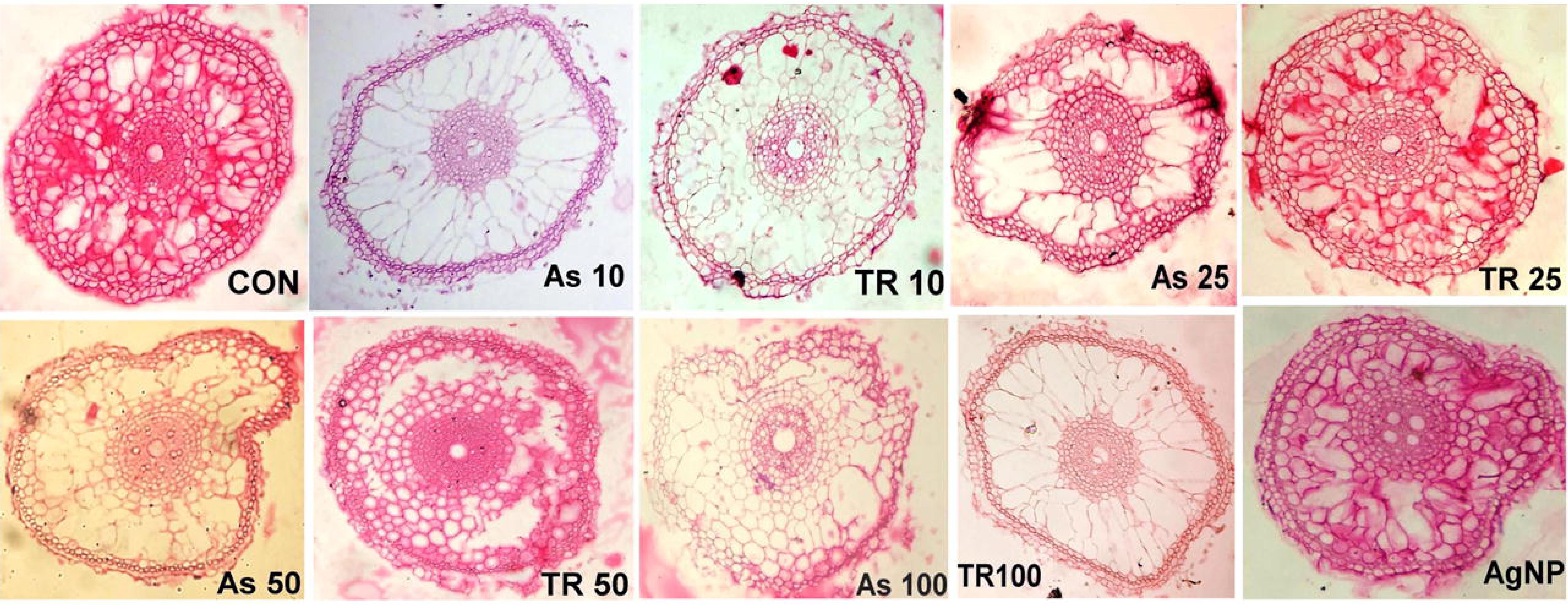
Cross section of rice roots showing anatomical changes upon exposure to all tested doses of As (III), TR and AgNP.

The scanning electron microscopic view revealed the formation of root hairs, smooth surface of root along with normal shaped root tips in CON, AgNP and all concentrations of TR; but in As exposed seedlings cracking in the root surfaces and root tips were visible (Fig.4). With increasing As concentrations, the formation of root hairs was also inhibited, which indicated the possible loss of cellular integrity as evinced by Nath et al. (2014) in *Oryza sativa*.

**Fig. 4.**
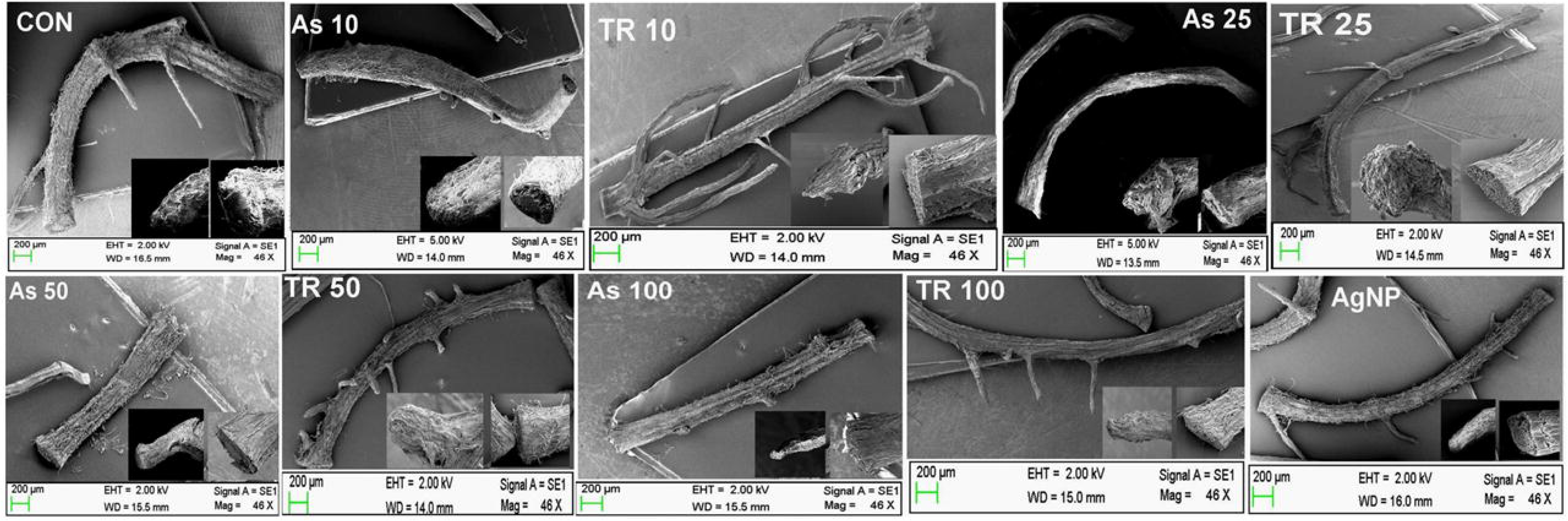
Scanning electron micrographs of roots after treatment at all tested concentrations of As (III), TR and AgNP.

### 3.3. Histochemical detection of ROS and superoxide anion (O^2-^)

*In situ* detection of arsenic, As(III) induced elevation of intracellular ROS generation and O^2-^ formation in root tips was assessed by fluorescent dye CM-H2DCFDA and DHE, respectively. CM-H2DCFDA diffuses passively into the cells, where its acetate groups are cleaved by esterases while the chloromethyl group reacts with intracellular glutathione and other thiols. Subsequent oxidation by H_2_O_2_ or other peroxides, it generates green fluorescence; another fluorescent dye DHE which is specific for superoxide anion (O^2-^) is oxidized by ROS to yield red fluorescence (Ghosh et al., 2016). A group of authors in their earlier reports showed the heavy metal induced ROS accumulation in roots of different plant species (Rao et al., 2011; Pandey et al., 2016). In the present study, both ROS and O^2-^ exhibited increasing pattern of fluorescent density with the increase in As concentrations; whereas, all respective TR doses and AgNP caused a significant reduction in the level of ROS generation and O^2-^ formation by showing almost similar fluorescent intensity with CON root tips (Fig.5 and 6) The results clearly indicated the application of nano pretreated (TR) water supplementation alleviate the As induced ROS accumulation and oxidative impairment in the roots of rice seedlings.

**Fig. 5.**
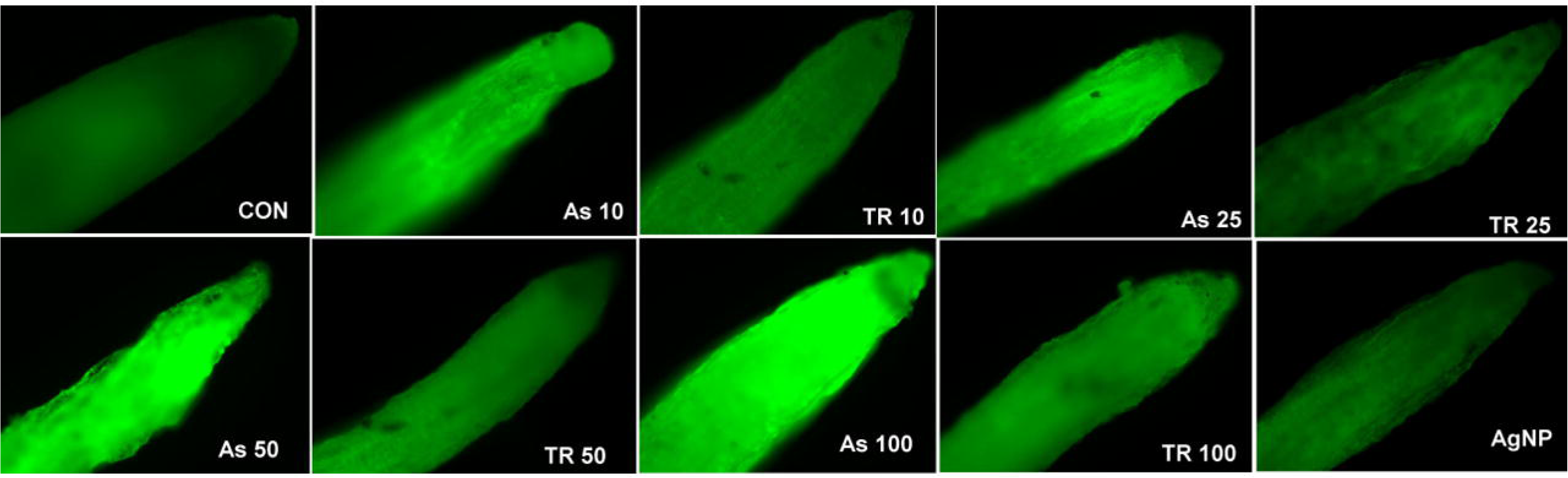
CM-H2DCFDA images of ROS formation in root tips in rice seedlings (MTU 7029) exposed to different As (III), TR and AgNP concentrations.

**Fig. 6.**
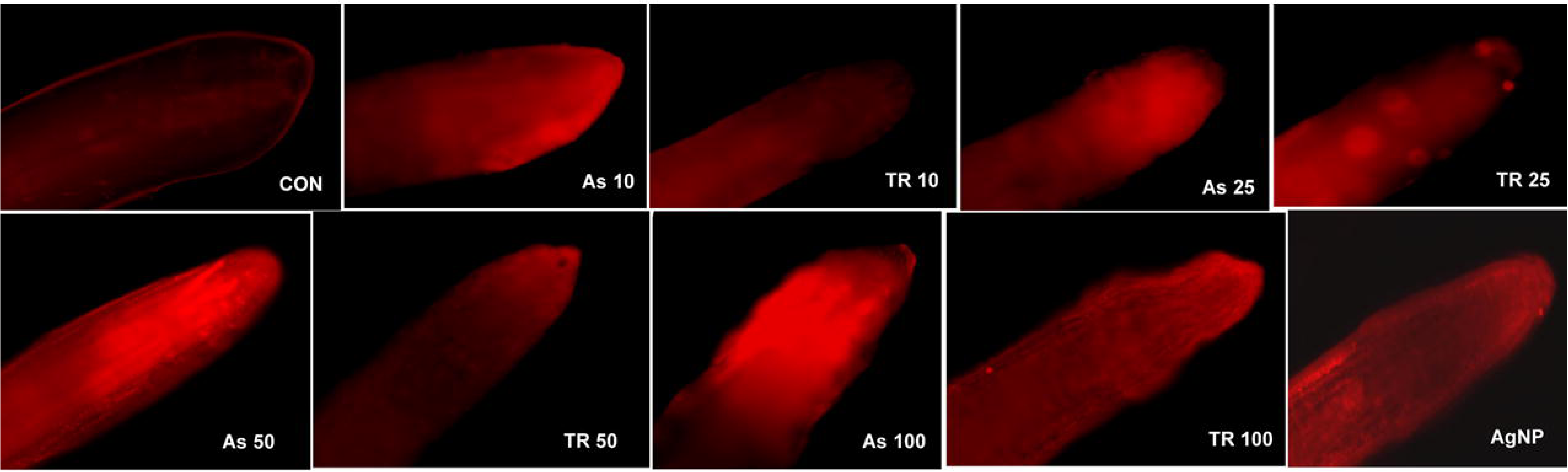
DHE images of ROS formation in root tips in rice seedlings (MTU 7029) exposed to all of the tested As (III), TR and AgNP concentrations.

**Fig. 7.**
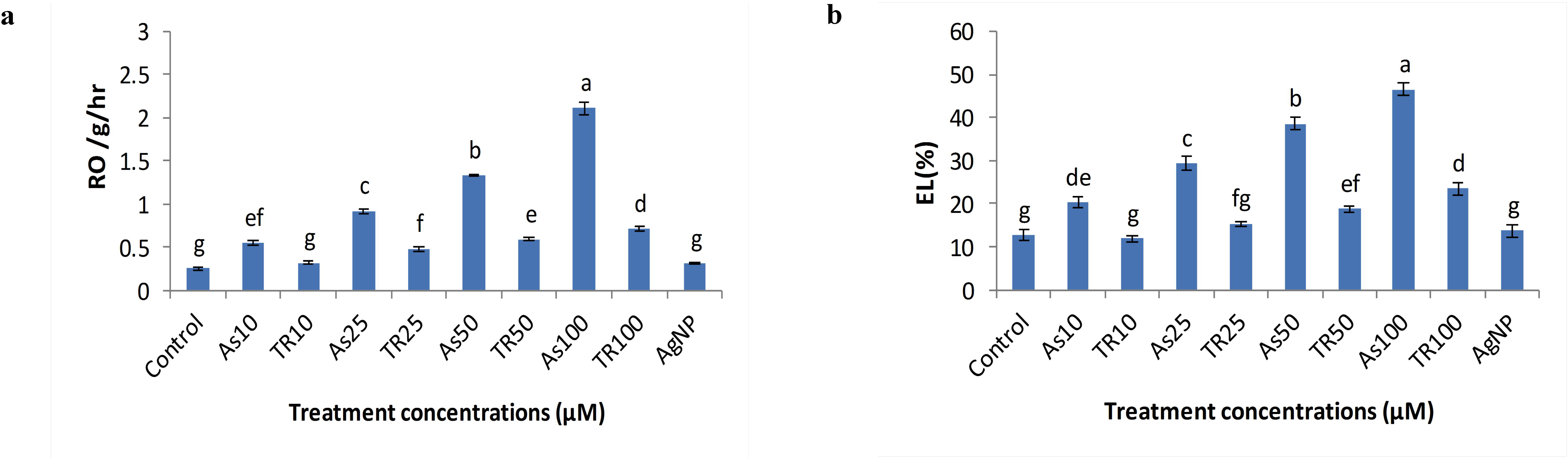
Response of rice roots to root oxidizabilty(a) and electrolyte leakage(b) after treatment at all of the tested concentrations. Data are expressed as means± S.E. of three replicate experiments. Vertical bars indicate standard errors. Mean values in columns followed by the different letters are significantly different at p ≤0.05 level according to Duncan multiple range test (DMRT).

**Fig. 8.**
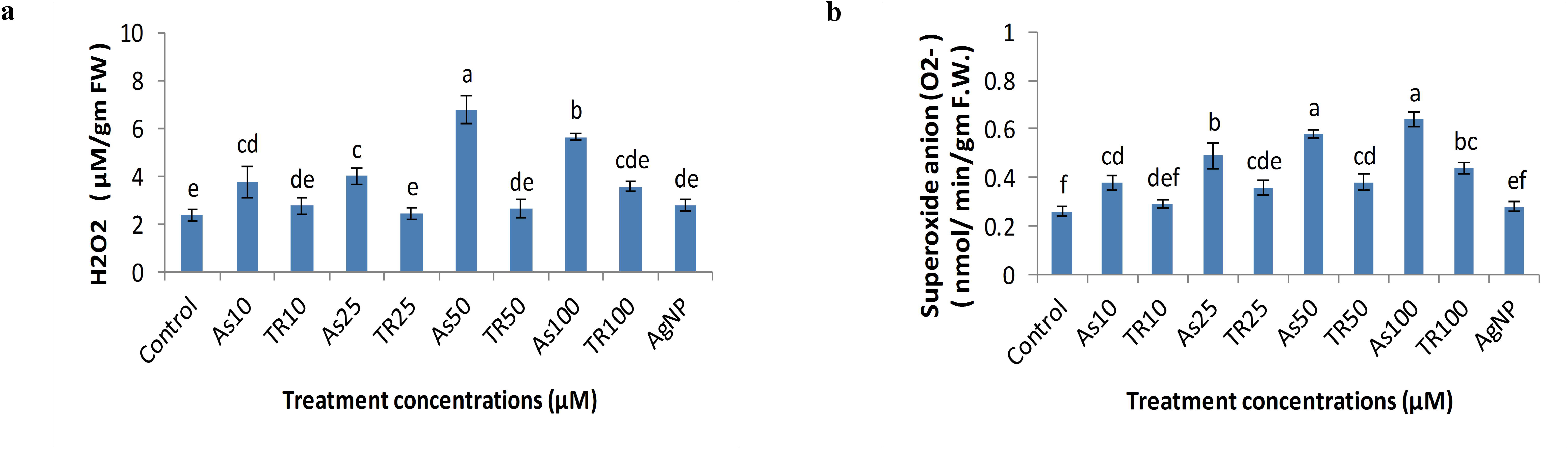
Effect of hydrogen peroxide content (a) and superoxide anion (b) on rice roots after treatment at different As (III), TR and AgNP doses. Data are expressed as means± S.E. of three replicated experiments. Vertical bars indicate standard errors. Mean values in columns followed by the different letters are significantly different at p ≤0.05 level according to Duncan multiple range test (DMRT).

### 3.4. Changes in root oxidizabilty (RO) and electrolyte leakage (EL%)

Root oxidizability increased significantly under As(III) exposure in hydroponic culture medium. With increasing As concentration RO gradually increased (Fig.7a), where maximum RO was recorded at 100 µM (∼10fold). Increased RO indicate a substantial diffusion of oxygen from the roots (Singh et al., 2007). Such enhancement of RO was further reduced with respective As concentration in the hydroponic culture medium amended with nano pretreated (TR) water. In all TR concentrations the level RO was reduced and in AgNP, it almost reached to the level of CON.

Arsenic exposure caused an electrolyte leakage (EL) from roots. Significant increase in (EL%) was observed in all concentrations of As, where the maximum leakage (∼4-fold increase) was found to be 46.53% at 100 µM of As in respective to control (CON). Subsequent application of nano-pretreated (TR) water supplementation with respective As concentration in the hydroponic culture medium resulted in a decrease of EL% and positive control AgNP resemblance with control (Fig.7b).

### 3.5. Alterations in hydrogen peroxide and superoxide anion content

Arsenic induced oxidative stress in rice plant (MTU 7029) was measured in terms of hydrogen peroxide (H_2_O_2_) and superoxide anion (O^2-^) content. Arsenic exposure significantly altered the H_2_O_2_ content in compare to control (CON). H_2_O_2_ content was increased in all of the As (III) concentrations where maximum content was exhibited at As50µM (Fig.8a). Notably, after maximum increase (∼3fold) at As50µM, a slight decline at As100 µM was observed. High H_2_O_2_ content in plants exhibited the toxic effect in terms of oxidative damage (Mishra et al., 2011, Singh et al.,2018). If such content is reduced, plants express better antioxidative defense system. In the present work, supplementation of nano pretreated (TR) water in hydroponic culture medium reduced the H_2_O_2_ content with respect to their As concentrations and the positive control AgNP were approximately similar to the control (CON) in H_2_O_2_ content.

Similar pattern of results was observed in case of superoxide anion (O^2-^) content where O^2-^ content was gradually increased with increasing As concentrations and maximum increase (∼2.5fold) was noticed at As100µM compared to control (Fig.8b). Enhanced O^2-^ generation upon arsenic exposure was observed in many plant species (Mishra et al., 2011). Superoxide anion is itself very destructive for plant cell as it causes oxidative damage to the cells and also serves as a source of other ROS in the cells (Halliwell and Gutteridge, 1984). Hence, plants need low O^2-^ content for betterment of their growth and metabolism. In the present study, the lowering of O^2-^ content was possible by the application of nano-pretreated (TR) water supplementation with the respective dose of As in hydroponic culture medium. The reduction at all concentrations of TR was not as lower as control (CON), but much lower as compared to their respective As doses (Fig.8b). In case of AgNP, the value was nearer to the control (CON) that strongly indicated nontoxic nature of AgNP to the rice seedlings.

### 3.6. Effect on proline and MDA content

Proline, a non-enzymatic antioxidant acts as a cytoplasmic osmoticum and also a good scavenger of free radicals (Dave et al., 2013). In the present study the level of proline content increased significantly in the rice seedlings upon As (III) exposure in hydroponic culture medium. The content increased in the leaf tissues in a concentration dependent manner upon As (III) exposure with the maximum increase (∼3.7 fold) being at As100 µM (Fig.9 a). The increase in free proline content upon As exposure has been observed earlier by several authors in different plants (Dave et al., 2013; Rahman et al., 2015; Pandey et al., 2016). Such enhancement of proline content was further reduced with respective As concentration in the hydroponic culture medium amended with nano pretreated (TR) water. In all TR concentrations the level proline content was reduced, while in TR10 and AgNP it almost reached to the level of CON.

**Fig. 9.**
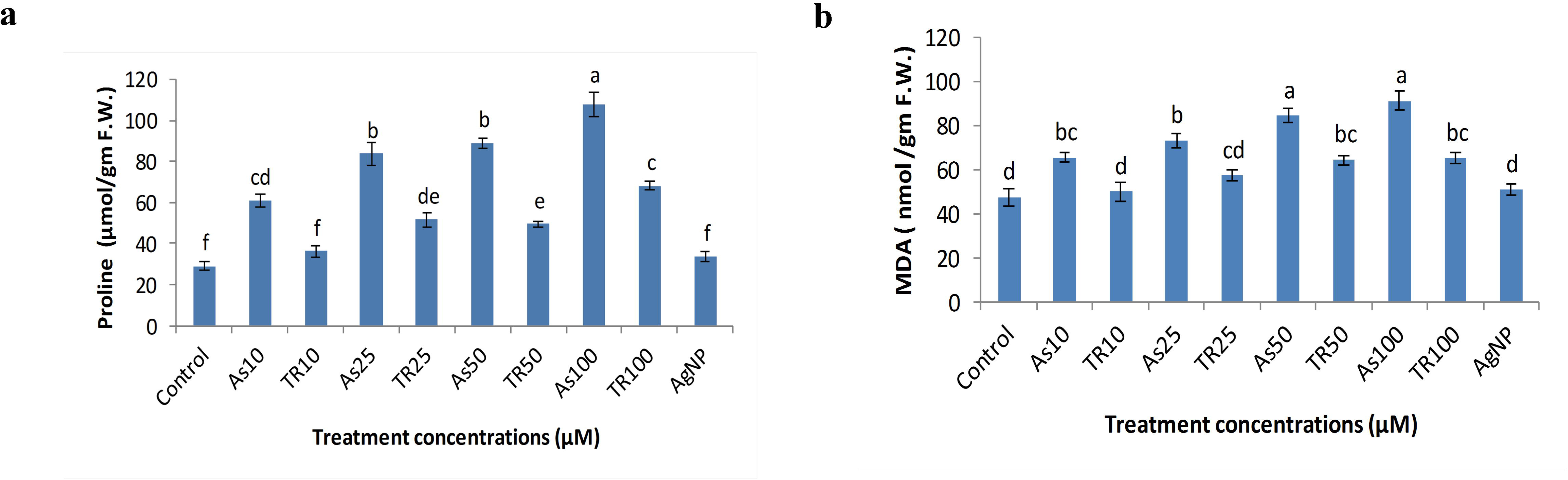
Effect of proline (a) and MDA (b) content to different tested concentrations of As (III), TR and AgNP. Data are expressed as means± S.E. of three replicated experiments. Vertical bars indicate standard errors. Mean values in columns followed by the different letters are significantly different at p ≤0.05 level according to Duncan multiple range test (DMRT).

MDA content also increased progressively in the leaf tissues of rice seedlings in a concentration dependent manner upon As (III) exposure with the maximum increase (∼2 fold) at As100 µM (Fig.9 b). Increased MDA content might be the reason for excess ROS production leading to the membrane damage due to peroxidation of poly unsaturated lipid (Singh et al., 2007). Our results were in accordance with the earlier reports by various authors under As stress (Pandey et al., 2016; Upadhyay et al., 2016). In contrast, increase level of MDA content in leaf tissues was further reduced with respective As concentration in the hydroponic culture medium supplemented with nano pretreated (TR) water. The reduction was evident in all of the TR concentrations, while in TR10 and AgNP it almost reached to the level of CON. This observation confirmed the nontoxic nature of this nanoparticle.

### 3.7. Effect on different antioxidant enzymes activity

Antioxidant enzymes are considered to be the indispensible components of plant defense system against metal induced oxidative stress (Weckx and Clijsters, 1996). Among the enzymatic antioxidants, SOD constitutes the first line of defense by dismutating the superoxide radicals, a major ROS created by the oxidative stresses (Gusman et al., 2013); three isoforms, Cu/zn-SOD, Mn-SOD and Fe-SOD have been found to exist in plants and are mostly nuclear encoded (Gill and Tuteja,2010). Another important ROS, H_2_O_2_ is generally scavenged by the action of another popular antioxidant enzymes,CAT and APX (Karuppanapandian et al., 2011). APX functions as a central component of ascorbate-glutathione cycle and more potential enzyme for controlling the intracellular ROS by reducing H_2_O_2_ into water (Upadhyay et al., 2016). Moreover, GR plays a crucial role by maintaining the reduced (GSH)/oxidized (GSSG) ratio in favor of ascorbate reduction (Noctor et al., 2002).

Plants exhibited tremendous variation in their antioxidant responses during As toxicity. In the present study, the spectrophotometric results revealed the differential modulation of antioxidant enzymes activities in both leaves and roots of rice seedlings (MTU 7029). The activity of H_2_O_2_ scavenging enzyme CAT was found to be lower in all concentration of As(III) treatment on both leaf and root, but in a differential manner. The maximum reduction in compare to control (CON) was noticed at As25 µM in leaf (∼1.5 fold), whereas in root same fold reduction ((∼1.5 fold) was noted at both As25 µM and As100 µM (Fig. 10 a, b).Hence, the low CAT activity might be responsible for higher accumulation of H_2_O_2_ in rice seedlings. This result was positively correlated with our findings where the increased level of H_2_O_2_ content was observed in all tested concentration of As(III) treatment in rice root. CAT activity was further enhanced in all of the TR concentrations as compared to the respective As concentrations tested alone. The activity in AgNP reached towards the control (CON), indicated the positive impact of this nanoparticle in the rice seedlings.

**Fig. 10.**
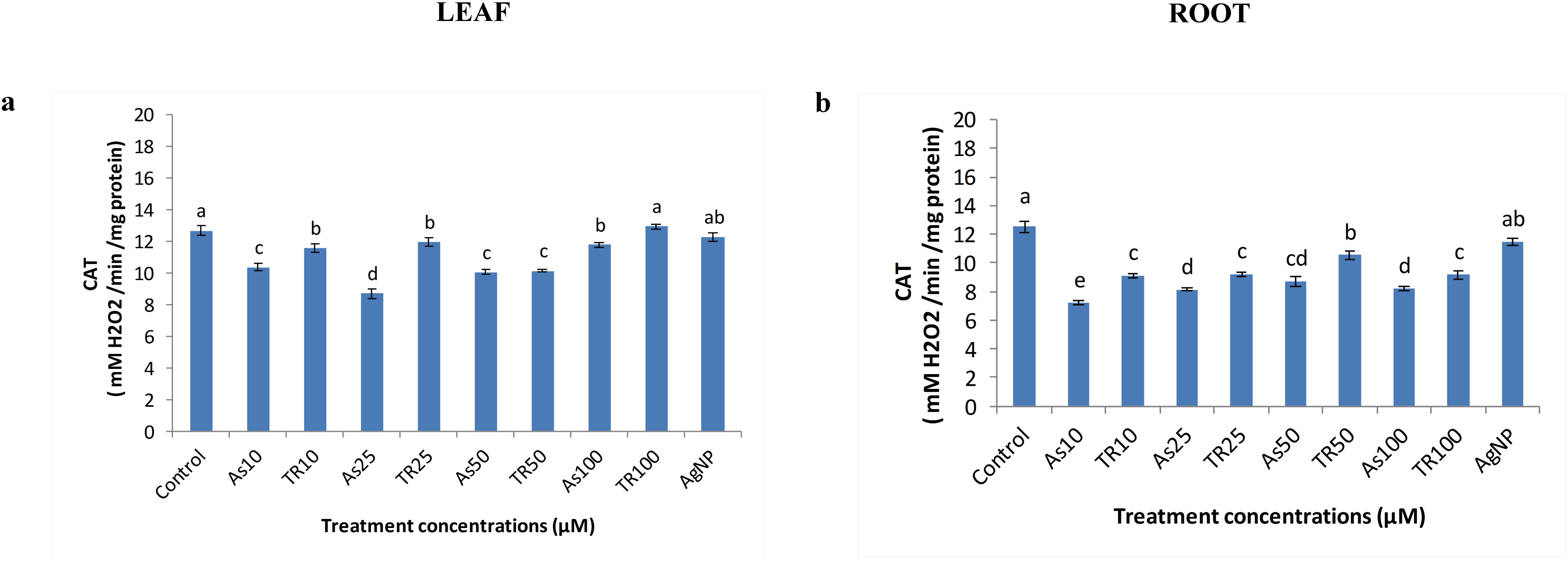
Differential response of catalase (CAT) activity in leaf (a) and root (b) after exposure to different concentrations of As (III), TR and AgNP. Data are expressed as means± S.E. of three replicated experiments. Vertical bars indicate standard errors. Mean values in columns followed by the different letters are significantly different at p ≤0.05 level according to Duncan multiple range test (DMRT).

The similar pattern of activity was also evinced in case of APX as compared to CAT activity. Like CAT, the activity of APX was also reduced after As (III) treatment. The activity pattern was different in leaf compared to root. In leaf the changes in the reduction of APX activity was noticed upto As25 µM, afterward in As100 µM no significant reduction was observed. Whereas, in root the response of APX activity was quite prominent; the significant reduction was noted in all concentration of As(III) treatment. The maximum reduction was recorded at As50 µM (∼1.6 fold of reduction) as compared to control (CON) (Fig.11 a,b). The decreased activity of APX under As stress might be either of low availability of reduced ascorbate, or by excess production of H_2_O_2_ (Hiner et al, 2000; Mallick et al, 2014). This reduced APX activity induced the overproduction of H_2_O_2_ and ROS that ultimately led to the retardation of growth and metabolism in rice seedlings(MTU 7029).Interestingly, upon supplementation with nano-pretreated (TR) water the activity of APX was further enhanced with their respective dose of As(III) treatment in hydroponic culture medium. The enhancement reached nearer to control (CON) in case of two TR concentrations (TR10 µM and TR 25 µM) also in the positive control AgNP.

**Fig. 11.**
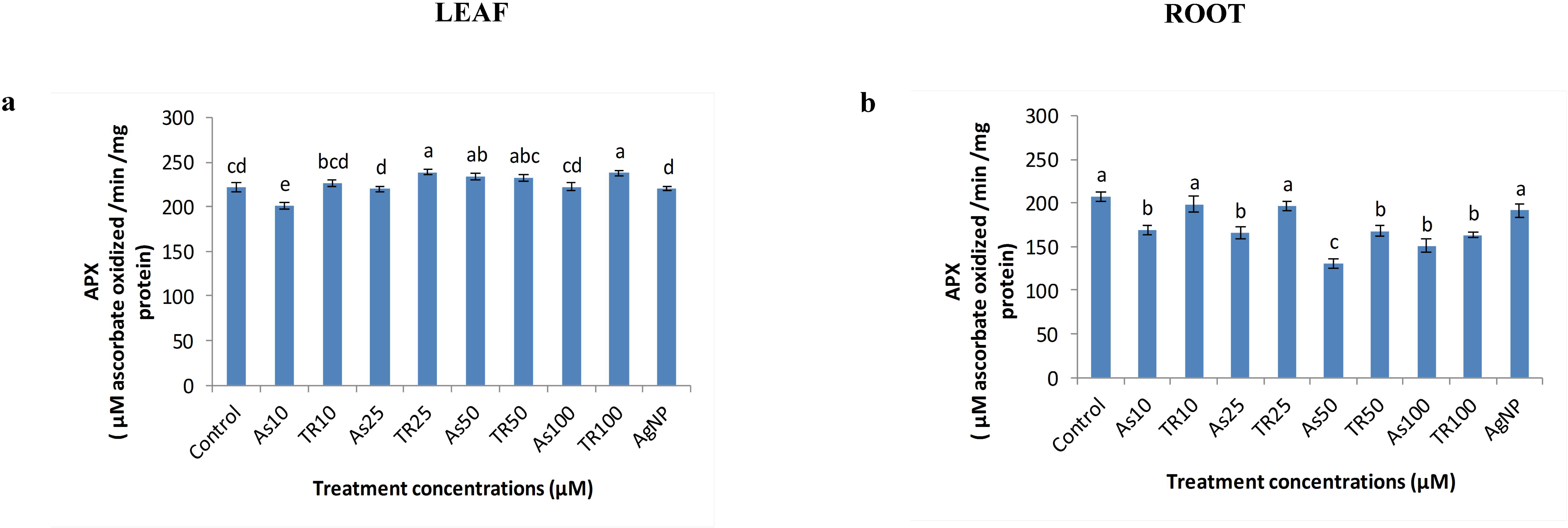
Differential response of ascorbate peroxidase (APX) activity in leaf (a) and root (b) after exposure to different concentrations of As (III), TR and AgNP. Data are expressed as means± S.E. of three replicated experiments. Vertical bars indicate standard errors. Mean values in columns followed by the different letters are significantly different at p ≤0.05 level according to Duncan multiple range test (DMRT).

**Fig. 12.**
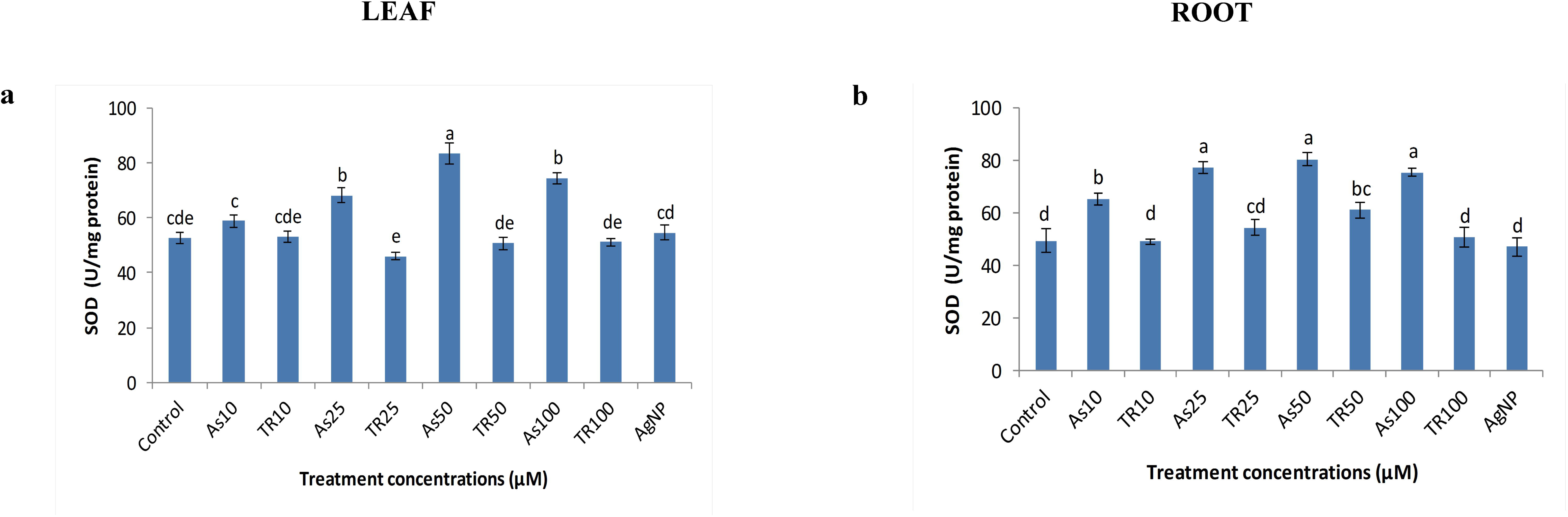
Differential changes in superoxide dismutase (SOD) activity in leaf (a) and root (b) after exposure to different concentrations of As(III),TR and AgNP. Data are expressed as means± S.E. of three replicated experiments. Vertical bars indicate standard errors. Mean values in columns followed by the different letters are significantly different at p ≤0.05 level according to Duncan multiple range test (DMRT).

**Fig. 13.**
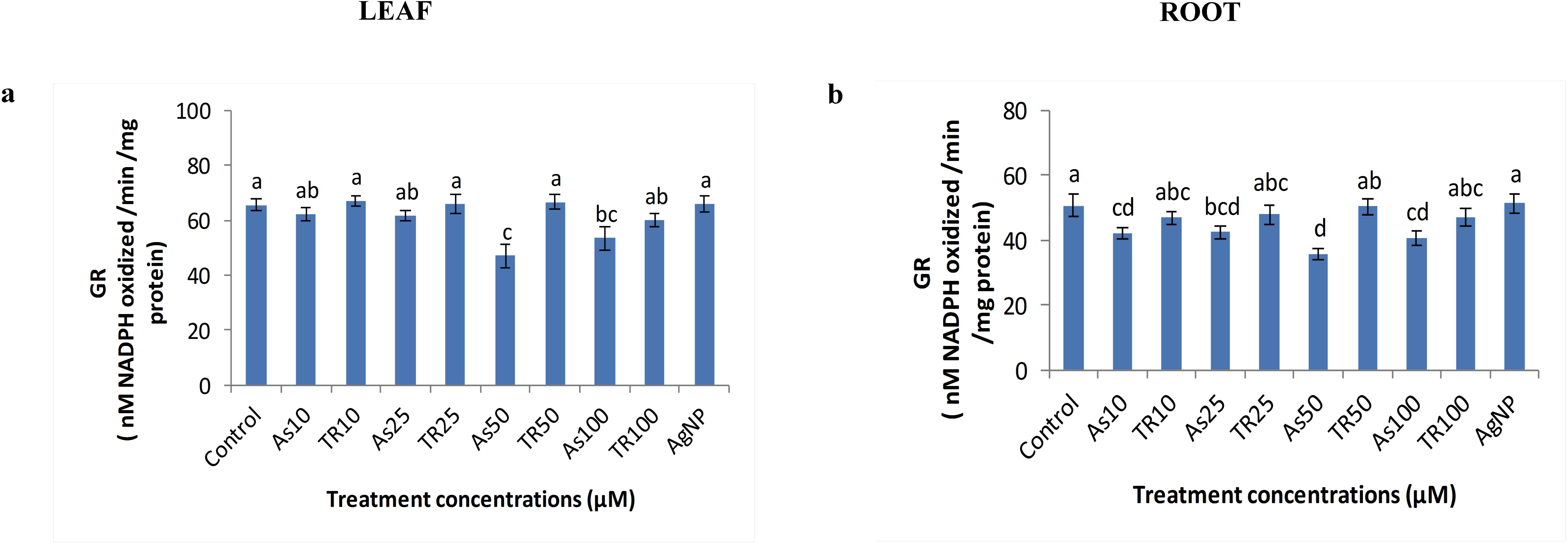
Differential activity of glutathione reductase (GR) in leaf (a) and root (b) after exposure to different concentrations of As (III),TR and AgNP. Data are expressed as means± S.E. of three replicated experiments. Vertical bars indicate standard errors. Mean values in columns followed by the different letters are significantly different at p ≤0.05 level according to Duncan multiple range test (DMRT).

Another two antioxidant enzymes tested so far in the present study were SOD and GR which also exhibited significant responses. In case of SOD, the activity was gradually increased with increasing As (III) concentrations up-to As50µM in both leaves and roots of rice seedlings. Notably, after maximum increase with an average of ∼1.6 fold in both leaf and root at As50µM, a slight decline was observed at As100µM, but that was also higher in respect of control (CON) (Fig.12a,b). Increased SOD activity might also be related to the higher accumulation of H_2_O_2_ as evident from the earlier studies on different plant system upon As stress (Shri et al., 2009; Mallick et al., 2014; Singh et al., 2018). Such increased activity of SOD was reduced upon supplementation with nano-pretreated (TR) water with their respective dose of As (III) treatment in hydroponic culture medium. The maximum fold of reduction was noted at TR 25 µM in case of leaf, while at TR10 µM and TR100 µM in root ((Fig.12a, b). In these particular concentrations of TR and AgNP the activity was looked similar to control (CON).

GR revealed the decreasing pattern of activity in both leaves and roots of rice seedlings upon As (III) treatment at different concentrations, but in a differential manner. In leaf the significant reduction was started after As25 µM while in root the reduction was noted from As10 µM (Fig.13a,b). The maximum loss of GR activity was found at As50 µM in both cases (∼1.4fold reduction in leaf and ∼1.5fold reduction in root). Decreased GR activity upon As treatment hampered the regeneration of glutathione and ascorbate to their reduced form in ascorbate-glutathione cycle, as reported by many authors in different plants (Tripathi et al., 2016; Jalil et al., 2023). Such decreased activity was further increased when the seedlings were supplemented with nano-pretreated (TR) water corresponding to their As concentrations tested alone (Fig.13a, b). The activity was noted approximately similar to the control (CON) in case of AgNP in both leaf and root of rice seedlings.

## 4. Conclusions

The present study deals with an efficient amelioration of arsenic induced toxicity by supplementation of nano-pretreated water in hydroponic culture medium of rice seedlings (MTU 7029). As (III) exposure in the growth medium disrupted the seedlings growth, physiological conditions as well as the antioxidant defense system by overproducing ROS and superoxide anions. Arsenic induced damage in the seedlings increased with increasing As(III) concentrations (10 μM to 100 μM) in the hydroponic growth medium; however, As50μM was supposed to be the most deteriorating concentration in many cases. Improvement of such deterioration was possible with the supplementation of nano-pretreated water, that indicated improved level of CAT, APX, SOD, GR and reduced ROS and superoxide anions formation as revealed by CM-H2DCFDA and DHE staining of the roots. Re-establishment of the redox status was also positively correlated with the other parameters examined in the present study. These included the betterment in the seedlings growth by enhancing root and shoot length, restoration of normal root morphology and anatomy as indicated by SEM and histology; decreased level of root oxidizability, electrolyte leakage percentage, H_2_O_2,_ MDA and proline content. Hence, the present findings could be useful for the farmer to apply this nano-pretreated water as a beneficial alternative of arsenic contaminated irrigation water in the cultivated land. Future investigations should include the large-scale implementation of this technology in both laboratory and field settings to gain further insights into the impact of this nanotechnology towards arsenic stress in the agricultural fields.

## Acknowledgements

The authors are thankful to University of Kalyani for providing their partial financial support through Personal Research Grant, DST-PURSE and DST FIST program. Authors are also thankful to Bose Institute for providing research support.

